# 17α-Estradiol Confers Limited Protection Against *APOE4* Phenotypes in Middle-Aged Female Mice

**DOI:** 10.64898/2026.08.06.743074

**Authors:** Cassandra J. McGill, Amy Christensen, Sara Namvari, Max A. Thorwald, Herbert Anson, Marc Vermulst, Caleb E. Finch, Bérénice A. Benayoun, Christian J Pike

## Abstract

Longevity-promoting interventions represent a promising strategy to mitigate brain aging and reduce Alzheimer’s disease (AD) risk. The NIA Interventions Testing Program identified the weak estrogen 17α-estradiol (17αE2) as a compound that extends healthspan and lifespan in mice, with effects observed primarily in males. Our recent work demonstrated that 17αE2 healthspan benefits were modulated by human apolipoprotein E (*APOE*) genotype such that aging phenotypes were improved more strongly in middle-aged male mice with targeted-replacement of the AD-associated *APOE4* allele compared to *APOE3*, the risk neutral and most common *APOE* allele. Here, we tested whether *APOE*-dependent, AD-relevant benefits of 17αE2 observed in males extend to females. Specifically, we treated 12-month-old *APOE3* and *APOE4* targeted-replacement female mice for 6 months with chow containing 0 or 14.4ppm 17αE2. We find that relative to *APOE3*, *APOE4* genotype largely exhibits more robust systemic phenotypes associated with aging, including increased adiposity, impaired glucose tolerance, and reduced energy expenditure. Further, we observe that treatment with 17αE2 yields modest improvements in some outcomes, including decreased adiposity and increased lean mass, glucose tolerance, and energy expenditure, though significant benefits are found only in *APOE4* females. In the CNS, we observed mixed effects of *APOE* genotype on behavioral performance and indices of brain aging, with *APOE4* females performing worse in the Barnes Maze and having higher levels of the AD-related peptide soluble β-amyloid, but no *APOE* genotype differences in cortical lipid raft oxidative damage. In contrast to its systemic effects, 17αE2 did not significantly improve neural outcomes in *APOE3* or *APOE4* females. These findings address the impact of biological sex on established protective effects of a longevity-promoting intervention against *APOE4* phenotypes, which have significant relevance to the prevention of age-related conditions including metabolic dysfunction, cognitive impairment and vulnerability to AD.

## Introduction

*APOE4* genotype strongly associates with the systemic and neural phenotypes linked to cognitive impairment and AD susceptibility [1–4]. The human population contains three *APOE* alleles (ε2, ε3, ε4), which differ by two single nucleotide polymorphisms causing amino acid substitutions at positions 112 and 158 of the protein [5, 6]. Longevity and vulnerabilities to both systemic (e.g., metabolic dysfunction) and neural (e.g., cognitive decline, AD risk) age-related outcomes are significantly affected by *APOE* genotype, with *APOE2* associated with increased longevity and reduced neural risks and *APOE4* with decreased longevity and elevated neural risks [4, 7, 8] relative to the most prevalent *APOE3* allele. One intriguing possibility is that the relationships among *APOE* genotype, mortality, and susceptibility to age-related impairments and diseases may be mediated in part by a broader impact of *APOE* on aging processes, with *APOE4* associating with progeroid phenotypes both systemically [1, 9–12] and neurally [4, 13–15] in humans and mice [2, 3, 16–18].

Given the overlaps among *APOE4*, aging, metabolism, and inflammation, longevity-promoting interventions present a novel way to combat *APOE4*-related aging and AD phenotypes. The NIA Interventions Testing Program identified 17α-estradiol (17αE2) as a longevity intervention that extends lifespan in male mice with no significant effect in females [19–22]. Although 17αE2 is a diastereomer of the potent estrogen 17β-estradiol, it exhibits comparatively weak classic estrogenic effects [23]. Acting in part via estrogenic signaling pathways [21], 17αE2 improves systemic metabolic outcomes of aging in male mice, resulting in reduced adiposity, improved glucose homeostasis, and decreased inflammation [19–21, 24]. Our group found that 17αE2 improves metabolic and neural outcomes in an *APOE*-dependent manner, with greater benefits observed in *APOE4* males [25], suggesting therapeutic potential of 17αE2 and perhaps other longevity interventions in reducing risks of *APOE4* genotype against age-related disorders.

Sex is a major biological variable influencing longevity, aging trajectories, and susceptibility to age-related diseases, including AD [26]. Females generally outlive males yet exhibit increased risk for AD, highlighting the importance of sex-specific mechanisms underlying aging and neurodegeneration. Further, the *APOE4* allele exhibits a significant sex bias, generally showing increased risks for indices of cognitive aging and AD in women [15, 27]. The metabolic and immune effects of 17αE2 in females remain largely unexplored. Given its established role in modulating *APOE4*-associated systemic and neural pathways, 17αE2 presents a promising pleiotropic intervention strategy for AD, with the potential to address both metabolic and inflammatory mechanisms linked to *APOE4* and aging. To investigate this possibility, we treated 12-month-old *APOE3* or *APOE4* targeted replacement female mice with a control or 17αE2-supplemented diet for six months. Our findings reveal that the *APOE4* genotype exacerbates metabolic and aging-related impairments, many of which are ameliorated by 17αE2 treatment. Notably, the benefits of 17αE2 were most pronounced in *APOE4* female mice, underscoring its potential as a strategy to reduce *APOE4*-driven metabolic and aging phenotypes and providing preclinical proof-of-concept for a personalized medicine approach.

## Methods

### Animals and treatment

All female mice were homozygous for knock-in of human *APOE3* or *APOE4*. The mice were generated from a breeding colony of EFAD (*APOE*^+/+^, *5xFAD*^+/-^) mice [28] but were non-carriers of the 5xFAD Alzheimer’s-related genes (*APOE*^+/+^, *5xFAD*^-/-^). The colony was started from breeding pairs generously provided by Drs. Mary Jo LaDu and Leon Tai (University of Illinois at Chicago). Mice were maintained in a vivarium under controlled temperature, a 12:12 light/dark schedule (lights on at 6:00am), group housing when applicable, and *ad libitum* access to water and food (except when specified otherwise). At 12.0-12.5 months of age, *APOE3* and *APOE4* mice were randomized to one of two dietary treatment groups (n=11-20/group): TestDiet 5LG6 chow [67.3% carbohydrate, 20.5% protein, 12.1% fat] formulated with 0 (Control diet) or 14.4 ppm 17αE2 (Steraloids, Newport, RI) by TestDiet (Richmond, IN). This 17αE2 dosage and delivery method were demonstrated to increase mean and maximum lifespan in male but not female UM-HET3 mice [20] and to exert modest estrogenic activity as indicated by increased uterine weight in ovariectomized female UM-HET3 mice [22]. We observed non-significant trends of elevated uterine weight in gonadally intact, 17αE2-treated female *APOE* mice, with the relationship more apparent in *APOE4* mice (**Supplementary Figure 1A**).

The experimental design is summarized in **Figure 1A**. In brief, the mice were maintained on the diets over a treatment period of 26 weeks, during which body weight and food consumption were recorded weekly. Body composition was measured at weeks 0 and 26 of treatment using a Bruker whole body composition analyzer (Bruker LF90 Minispec, Bruker Optics, Billerica, MA). Mice were monitored daily for overall health, appearance, and euthanasia criteria, including >20% body weight decrease, lethargy, and poor grooming. Of the 70 animals enrolled, 10 animals were euthanized before completing the study (3 *APOE3* Control, 3 *APOE3* 17αE2, 2 *APOE4* Control, 2 *APOE4* 17αE2). After the 26-week experimental period, mice were euthanized through inhalation of carbon dioxide after overnight food withdrawal, followed by transcardial perfusion with 20mL ice-cold 0.1M PBS. The brains were rapidly removed, and one hemibrain was immersion-fixed for 48 hours in 4% paraformaldehyde/ 0.1M PBS, then stored at 4°C in 0.1M PBS/ 0.03% NaN_3_ until processing for immunohistochemistry. The other hemibrain was dissected into cerebral cortex, hippocampus, and hypothalamus. All portions were snap-frozen for RNA or protein extraction. Plasma was collected and stored at −80°C. Visceral, retroperitoneal, and subcutaneous fat pads and livers were dissected and weighed; all were snap-frozen. All procedures were conducted in accordance with National Institutes of Health guidelines, under the supervision of veterinary staff, and following a protocol (#21269) approved by the University of Southern California Institutional Animal Care and Use Committee.

**Figure 1.**
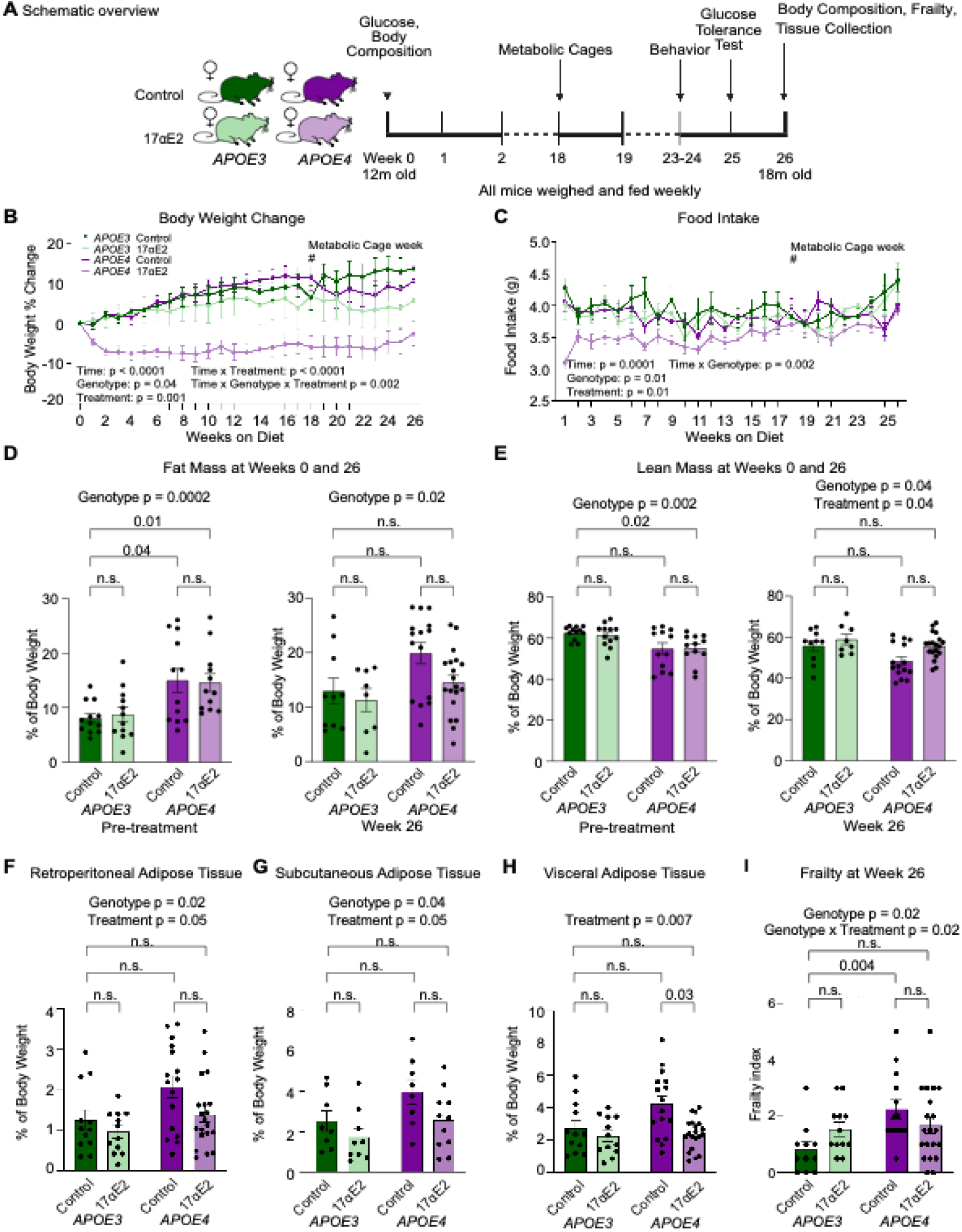
17α estradiol decreases body weight and food intake in aged female *APOE4* targeted replacement mice. (A) Schematic overview. Food intake and body weight were measured weekly. Fat/Lean mass was measured prior to treatment and after 26 weeks of treatment. (B) Percent body weight change across the 26 weeks of treatment (n=11-20/group). (C) Food intake (g) across the 26 weeks of treatment (n=11-20/group). (D) Fat mass measured by the Bruker Mini Spec (n=9-19/group). (E) Lean mass measured the Bruker Mini Spec (n=9-19/group). In (D) and (E), shaded panels indicate pre-treatment. (F) Percent of retroperiatoneal adipose tissue compared to overall body weight after 26 weeks of 0 or 14.4ppm 17αE2 (n=11-20/group). (G) Percent of subcutaneous adipose tissue compared to overall body weight after 26 weeks of 0 or 14.4ppm 17αE2 (n=8-11/group). (H) Percent of visceral adipose tissue compared to overall body weight after 26 weeks of 0 or 14.4ppm 17αE2 (n=11-20/group). (I) Measure of systemic aging using a phenotypic 26-point frailty index (n=11-20/group). In panels (B) and (C), dark green lines indicate *APOE3* control, light green lines indicate *APOE3* 17αE2, dark purple lines indicate *APOE4* control, and light purple lines indicate *APOE4* 17αE2. In (D-I), dark green indicates *APOE3* control, light green indicates *APOE3* 17αE2, dark purple indicates *APOE4* control, and light purple indicates *APOE4* 17αE2. Data show mean ± SEM.

### Liver oil red O (ORO) staining and quantification

Frozen livers were sectioned at 13 µm using a cryostat at −14°C. Four sections were collected on the same glass slide for each animal (N=8/group). The slides were stored at −20°C until ORO staining, which was performed as previously described [25]. In brief, frozen liver sections were brought to room temperature, dipped 10 times into freshly prepared 60% triethyl phosphate, and then stained with a solution of 0.5% Oil Red O/60% triethyl phosphate for 16 minutes. Sections were rinsed in a gentle stream of running water for 2 minutes and put into clean water before mounting with pre-warmed glycerin jelly. Stained sections were stored at room temperature for one day, after which high-magnification brightfield images (four sections/animal, two fields/section, 40X oil objective) were collected by unbiased sampling for a total of 8 liver images per animal. Images were captured using an Olympus BX50 microscope and DP74 camera paired with a computer running CellSens software v1.11(Olympus). Images were converted to grayscale and thresholded using NIH ImageJ 1.50i to yield binary images separating positive and negative staining. The Analyze-Measure tool was used to obtain values representing the percentages of the ORO-positive area. For each animal, values were averaged across all 8 images.

### Blood glucose measurements

At Week 0, mice were fasted for 16 hours overnight after which blood glucose levels were measured. At Week 25, mice were assessed using a glucose tolerance test (GTT) following 16 hours of fasting. Baseline blood glucose was determined, and then animals were orally gavaged with 20% D-glucose in water (2g/kg). Blood glucose levels were measured through tail vein bleed and recorded 15, 30, 60, and 120 minutes after administration of the glucose using the Precision Xtra Blood Glucose and Ketone Monitoring System (9881465; Abbott Laboratories, Abbott Park, IL).

### Metabolic profiling

Metabolic profiling was performed using the TSE PhenoMaster System for fully automated assessment of home cage metabolic monitoring of food and water intake (including amount and time patterns), energy expenditure (by indirect calorimetry), and voluntary locomotor activity (infrared x-y-z sensors for documentation of movements in 3-dimensions). Animals were recorded for 5 days, and the final two full days (midnight to 11:59pm) were used for analysis to allow the animals to habituate for 3 days prior to data collection. Raw measurements were aggregated into hourly means for each animal to allow comparison across the circadian cycle. For phase-specific analyses, measurements were averaged across the light phase (06:01–17:59), dark phase (18:00–06:00), and the full 48-hour period. Linear regression models were used to evaluate the effects of experimental group while adjusting for body weight. Separate models were fit for measurements averaged across the full day, light phase, and dark phase. Estimated marginal means were calculated using the emmeans v.2.0.0 package in R, and pairwise group comparisons were performed using Tukey-adjusted tests. Results are reported as estimated marginal means ± standard error unless otherwise indicated.

### Visceral adipose tissue RNA isolation and RNA-seq library preparation

For RNA isolation, an approximately 15mm in diameter amount of visceral adipose tissue was resuspended in 1 ml of Trizol reagent (15596018; Invitrogen, Carlsbad, CA) and homogenized using 20 strokes with a dounce homogenizer. The manufacturer’s protocol was followed except for an additional 10-minute 12,000 x *g* spin prior to the addition of chloroform, a step that reduces lipid content. The resultant RNA pellet was treated with RNase-free DNase I (Epicentre) for 30 minutes at 37°C, and a phenol/chloroform extraction was performed to isolate RNA. RNA quality was assessed with the Agilent TapeStation platform using the RNA integrity number before sending to Novogene Corporation (Sacramento, CA) for library preparation and sequencing. Paired-end 150-bp reads were generated on the Illumina NovaSeq6000 platform at the Novogene Corporation (Sacramento, CA).

### RNA-seq bioinformatic analysis pipeline

Paired-end 150-bp reads were hard-trimmed to 141 bp using Trimmomatic v0.39 [29]. Trimmed reads were mapped to the mm39 genome reference using STAR v.2.7.0e [30]. Read counts were assigned to genes from the UCSC mm39 reference using subread v.2.0.2 [31] and were imported into R version 1.4.1717 to perform differential gene expression analysis.

Only genes with mapped reads in at least half of the RNA-seq libraries were considered expressed and retained for downstream analysis. We used surrogate variable analysis (sva) to estimate and correct for unwanted experimental noise [32]. R package ‘sva’ v.3.56 was used to estimate surrogate variables and the removeBatchEffect function from ‘limma’ v.3.64.3 was used to regress out the effects of surrogate variables from raw read counts. The ‘DESeq2’ R package (DESeq2 v.1.48.2) was used for further processing of the RNA-seq data in R [33]. Genes with FDR < 5% were considered statistically significant and are reported in **Supplementary Table 1**. We found a non-linear relationship of treatment between the genotypes and thus modeled treatment and genotype separately. The following comparisons were performed: *APOE3* control to *APOE4* control, *APOE3* control to *APOE3* 17αE2, and *APOE4* control to *APOE4* 17αE2.

### Dimensionality reduction

To perform multi-dimensional scaling (MDS) analysis [34], we used a distance metric between samples based on the Spearman’s rank correlation value (1-Rho), which was then provided to the core R command ‘cmdscale’. Dimensionality reduction was applied to DESeq2 VST-normalized counts.

### Functional enrichment analysis

The Gene Set Enrichment Analysis (GSEA) paradigm through its implementation in the R package ‘clusterProfiler’ v4.16.0 [35], and Bioconductor annotation package ‘org.Mm.eg.db’ v3.21.0 were used to perform the functional enrichment analysis. The DEseq2 t-statistic was used to generate the ranked list of genes for functional enrichment analysis, for both genotype and treatment effects. All significant GO terms are reported in **Supplementary Table 1**.

### Behavioral assessments

#### Barnes maze

The Barnes maze test was performed at Week 23 using a modified Barnes maze protocol [36]. The maze consisted of an open circular platform (91.5cm in diameter) with 20 evenly spaced holes (5cm in diameter) located along the border with a rectangular escape box (11cm L x 5cm W x 5xm H) located beneath one hole. Using spatial-visual clues on each side of the platform, mice were trained to find the escape box. On each day of testing, mice were taken into the behavior room 30 minutes prior to testing to allow them to acclimate. On the first day of the Barnes maze, the escape box was removed, and mice were habituated to the maze. Each mouse was given a single habituation trial in which they were placed in an opaque cylinder in the center of the maze. After 10 seconds had elapsed, the cylinder was removed, allowing the mice to explore the maze freely for 3 minutes under red light. On the second day of the Barnes maze, mice were placed in a cylinder in the center of the maze, a bright light and buzzer were turned on, and after 10 seconds had elapsed, the cylinder was removed. Mice were gently guided into the escape box, after which the light and buzzer were turned off. Mice stayed in the escape box for 1 minute before they were returned to their home cages. After this initial training session, each testing day (including the initial training session day) consisted of 3 training trials per day with an intertrial interval of 15 minutes. Between trials, mice were kept in their own home cages, and the maze was cleaned with 70% ethanol. Acquisition training continued for 3 more days (4 training days total). During each trial, mice were placed in a cylinder in the center of the maze, the light and buzzer were activated, and after 10 seconds, the cylinder was removed. Mice were given 3 minutes to explore freely and locate the escape box. The trial ended when 3 minutes elapsed, or mice found and entered the escape box. Once a mouse found and entered the escape box, the light and buzzer were turned off, and the mouse stayed inside the box for 1 minute. If the mouse did not find the escape box after 3 minutes of exploration, they were gently guided into the escape box and stayed inside the box for 1 minute. To ensure that the hidden escape box was not visible to the mice, three decoy boxes (5cm L x 5cm W x 2.5cm H) were placed throughout the maze. 48 hours after the last acquisition trial, mice were given a probe trial in which the escape box was removed. On the probe day, mice were placed in the cylinder in the center of the maze, and a light and buzzer were turned on. After 10 seconds, the cylinder was removed, and mice were given 3 minutes to explore freely. All tests were recorded using Noldus Ethovision XT software version 14.

#### Open field

A standard open field test was performed at Week 24 on all mice. The mice were moved to the behavior room 30 minutes prior to testing to acclimate. Mice were then placed into an open field box (40cm x 40cm) and allowed to explore freely for 5 mins. The following behaviors were recorded using Noldus Ethovision XT software version 14: time in center (s); travel distance (cm); ambulatory time (s).

#### Novel object placement/recognition

Novel object placement (NOP) was performed at Week 24 of treatment using a modification of a previously described protocol [37]. Each day during the 3 days of assessment, mice were taken into the behavior room to habituate for 30 minutes prior to testing. Beginning 24 hours after the open field test, mice were placed in the empty arena (40cm x 40cm), facing the wall that was nearest to the experimenter, and explored for 5 minutes to habituate to the arena. Twenty-four hours after the first habituation, a second habituation was performed in the same way. Twenty-four hours after the second habituation session, the sampling trial consisted of two identical objects placed in the northeast and northwest corners of the arena, and mice were placed in the arena with their heads positioned opposite the objects. Mice were allowed to explore both objects for 5 minutes before being returned to their home cages. Fifteen minutes after sampling, mice were placed in the testing arena, where one of the identical objects was moved to the southeast or southwest corner (novel object placement). The location of the moved object was counter-balanced across all mice. Mice were allowed to explore both objects for 5 minutes before being returned to their home cage. Fifteen minutes after NOP, mice were placed in the testing arena in which one of the identical objects was replaced with a novel object. Mice were allowed to explore both objects for 5 minutes before being returned to their home cage. Data are presented as a discrimination index, which is defined as the time spent within 2cm of the novel object minus the time spent with the familiar object divided by total exploratory time. All tests were recorded using Noldus Ethovision XT software version 14.

### Amyloid-ß Measurements

Brain cerebral cortices were homogenized with a motorized pestle in RIPA buffer without SDS (30 mg tissue: 150 μL) with protease (P2714, Millipore, Bedford, MA, USA) and phosphatase (78427, Thermo Fisher Scientific, Waltham, MA) inhibitors. Homogenates were centrifuged at 10,000 x *g* for 1 hour at 4°C, and supernatant was recovered for evaluation of soluble amyloid-*ß* peptides. Protein concentration was determined using a BCA assay (A55864, Thermo Fisher Scientific, Waltham, MA). Supernatant containing 100 µg protein was added to each sample well (duplicate wells for each animal) of a MesoScale Diagnostics Amyloid (6E10) V-PLEX plate. Samples were incubated with detection antibody overnight at 4°C on a shaker. The plate was washed and read the following morning.

### Lipid Raft Oxidative Damage

Lipid rafts (LR) were isolated by kit (LR-039, Invent Biotechnologies, Plymouth, MN) using 35mg of cerebral cortex per animal. LRs have been previously validated against traditional ultracentrifugation methods [38]. Protein amounts were quantified using Pierce’s 660nm assay (22660, Thermo Fisher Scientific, Waltham, MA). Lipid raft protein lysates (5 µg/sample) were loaded onto a dot blot apparatus (Bio-Rad, Hercules, CA, USA) and filtered through 0.45μm PVDF membrane for 2 hours by gravity filtration. Membranes were stained with Revert 700 (926-11011, LICOR, Lincoln, NE, USA), imaged, and blocked for 1 hour with Intercept blocking buffer (927-70001, LICOR) before incubation for 16 hours with primary antibodies against 4-hydroxynonenal (4-HNE; 1:1000 dilution, ABN249, Millipore, Bedford, MA, USA) and nitrotyrosine (NT; 1:1000 dilution, 06-284, Millipore, Bedford, MA, USA). Membranes were visualized on a LICOR 9120 using fluorescent-conjugated secondary antibodies. Images were analyzed by ImageJ and corrected by total protein load.

### Frailty measurement

The frailty index (FI) score was calculated for each mouse using a 26-point frailty index, which was modified from a 31-item frailty index[39] to omit measures of vestibular disturbance, hearing loss, and vision loss due to the subjective nature of the scoring to ensure consistency across investigators, as previously described [25]. Other measures excluded from FI assessment include body weight and body composition due to baseline differences across *APOE* genotype that are measured separately, and body temperature due to prior studies showing no difference. FI assessment included evaluation of the integument, the physical/musculoskeletal system, the ocular/nasal system, the digestive/urogenital system, the respiratory system, body temperature, and signs of discomfort. The severity of each deficit was rated with a simple scale: 0, 0.5, and 1. A score of 0 was given if there were no signs of a deficit, a score of 0.5 was given to a mild deficit, and a score of 1 indicates a severe deficit. All these values were summed, giving a frailty score between 0 and 26 for each mouse. The researcher performing the measurement was blinded to all groups.

### DNA methylation sequencing and epigenetic age calculation

Frozen liver tissue (n=8/group) was processed by the Zymo Research DNAge Service (Tustin, CA). Briefly, DNA was purified from the frozen liver using the Quick-DNA Miniprep Plus kit (D4068, Zymo Research, Tustin, CA). After quality and quantity checks, bisulfite conversion was performed using the EZ DNA Methylation-Lightening kit (D5030, Zymo Research, Tustin, CA). Samples were enriched for >500 age-associated gene loci and sequenced on an Illumina NovaSeq6000 instrument. Sequenced reads identified by Illumina’s base calling software were aligned to the mouse reference genome using Bismark. Cytosine methylation level was determined as the number of reads reporting a C, divided by the total number of reads reporting a C or T. DNA methylation values were used to assess DNAge according to Zymo’s proprietary DNAge predictor.

### Statistics

All data are reported as the mean ± the standard error of the mean. Data were analyzed using GraphPad Prism version 5 (repeated-measure behavioral and metabolic data) or R version 4.5.2 (all other measures). Two-way repeated measures ANOVA, followed when appropriate by Tukey *post hoc* tests, was run for all data measured over time. For all other measures, the non-parametric Aligned Rank Transform ANOVA was performed. Comparisons with *p* < 0.05 were considered statistically significant.

## Results

### 17αE2 improves body composition in female APOE4 mice

The *APOE4* allele is associated with reduced lifespan in humans and mice [8]. To determine if the longevity-promoting intervention 17αE2 offers greater protection against age-associated systemic phenotypes in the context of the *APOE4* allele, we used strains of mice with humanized *APOE3* or *APOE4* sequences [28]. Our previous work in male *APOE* knock-in mice showed that 17αE2 had more pronounced effects on metabolic, neural, and frailty outcomes in *APOE4* carriers [25]. To determine if this also applied to females, *APOE3* and *APOE4* knock-in female mice were maintained on chow supplemented with 0 (Control) or 14.4ppm 17αE2, the dose previously shown to have anti-aging effects [22]. Treatment was initiated in early middle age (12 months of age) and maintained for a period of 6 months (**Figure 1A**). 17αE2 was associated with reductions in body weight across both genotypes, but with greater initial effect in the *APOE4* mice (**Figure 1B**). Consistent with this, there were only significant decreases in food intake in the *APOE4* mice in the initial weeks of treatment (**Figure 1C**). At week 0 of treatment, *APOE4* mice exhibited significantly higher body weight compared to *APOE3* mice (**Supplementary Figure 2A**), as well as significantly higher percent fat mass (**Figure 1D**) and lower percent lean mass (**Figure 1E**). Body composition analysis at week 26 revealed a similar pattern of 17αE2 treatment improvements, with only *APOE4* mice exhibiting a trend for reduced relative fat mass (**Figure 1D**) and significantly increased relative lean mass (**Figure 1E**). Importantly, following 17αE2 treatment, *APOE3* and *APOE4* mice did not significantly differ in measures of body morphometry. These findings were paralleled by tissue weights of retroperitoneal, subcutaneous, and visceral adipose depots, in which *APOE4* was associated both with higher adiposity under the Control diet and reductions in adiposity with 17αE2 treatment (**Figure 1F-H**). We assessed the ability of 17αE2 to impact aging phenotypes in middle-aged female *APOE* mice using a 25-point frailty index [39]. The frailty index measures visible markers of aging including the physical/musculoskeletal system, the ocular/nasal system, the respiratory system, the digestive/urogenital system, and observable signs of discomfort [39]. Control *APOE4* mice had a significantly higher frailty index compared to *APOE3* control, a finding consistent with observations in males [25]. There was a significant interaction between genotype and treatment, with a trend for higher frailty in 17αE2-treated *APOE3* mice than control, whereas 17αE2-treated *APOE4* mice displayed lower frailty compared to control (**Figure 1I**). Consistent with findings in genetically heterogeneous mice with wild-type murine *Apoe* [24], as well as male mice with human *APOE3* or *APOE4* [25], 17αE2 was associated with reduced food intake, body weight, adiposity, and frailty in female mice with knock-in of human *APOE4.* Conversely, *APOE3* animals showed modest, statistically nonsignificant changes in body weight, food intake, and adiposity with 17αE2 treatment, as well as a trend for increased frailty, suggesting genotype-specific mechanisms of 17αE2-induced actions.

### 17αE2 divergently impacts the visceral adipose tissue transcriptome in APOE3 and APOE4 mice

To investigate the molecular changes related to the observed metabolic outcomes, we performed bulk RNA-seq on the visceral adipose tissue (VAT) of control and 17αE2*-*treated *APOE* targeted replacement female mice (**Figure 2A**). The VAT was chosen for deeper analysis because it showed the greatest differences by both genotype and treatment (**Figure 1H**) and is an established regulator of functional metabolic outcomes [40, 41]. Multidimensional scaling analysis showed clear separation by both genotype and treatment (**Figure 2B**). Between control *APOE3* and *APOE4* groups, there were 1445 differentially expressed genes (DEGs), and Gene Set Enrichment Analysis (GSEA) using Gene Ontology terms showed *APOE4* control VAT was enriched for immune- and inflammatory-related terms, while *APOE3* control VAT was enriched for mitochondrial- and metabolism-related terms (**Figure 2C, Supplementary Table 1A,B**). There were 1401 DEGs between *APOE4* 17αE2 and *APOE3* control, suggesting treatment did not substantially change the *APOE4* transcriptome to be more similar to *APOE3* controls (**Figure 2D, Supplementary Table 1C,D**). Between control and treated *APOE3* VAT, there were 217 DEGs, with trends for immune-related term enrichment with 17αE2 and metabolism-related terms significantly enriched in controls (**Figure 2D, Supplementary Table 1E,F**). Between control and treated *APOE4* VAT, there were 254 DEGs, with methylation- and plasma lipoprotein remodeling terms enriched with 17αE2 and extracellular matrix-related terms enriched in controls (**Figure 2E**, **Supplementary Table 1G,H**). Twelve genes were differentially expressed (FDR < 10%) in the same direction with 17αE2 treatment in both genotypes (**Supplementary Figure 3A**, **Supplementary Table 1**). Among these genes was *Apoc2*, which regulates lipoprotein metabolism through activation of lipoprotein lipase. Interestingly, the highest enriched gene set with 17αE2 treatment in the *APOE4* comparison was *high-density lipoprotein particle* (**Figure 2G, Supplementary Table 2H**). *APOE3* and *APOE4* differ in their lipid-binding efficiency due to a structural variation at amino acid position 112, with *APOE3* being more efficient in binding lipids and stabilizing lipoproteins [42]. In contrast, *APOE4’s* structural rigidity reduces its lipid-binding efficiency, favoring interactions with very low-density lipoproteins (VLDL) and contributing to higher cholesterol levels and disease risk [42]. We assessed shared GO terms found in both the *APOE3* and *APOE4* comparisons to their treated counterparts (**Figure 2G, Supplementary Table 1J**). Contrary to 17αE2’s effect in *APOE4* VAT, *APOE3* treatment with 17αE2 was negatively enriched for lipoprotein-related terms. GO terms relating to mitochondria and actin were also divergently enriched between the genotypes, while inflammation-related terms were similarly positively enriched. While 17αE2 elicits transcriptional changes in both genotypes, phenotypically there are only benefits in *APOE4* mice.

**Figure 2.**
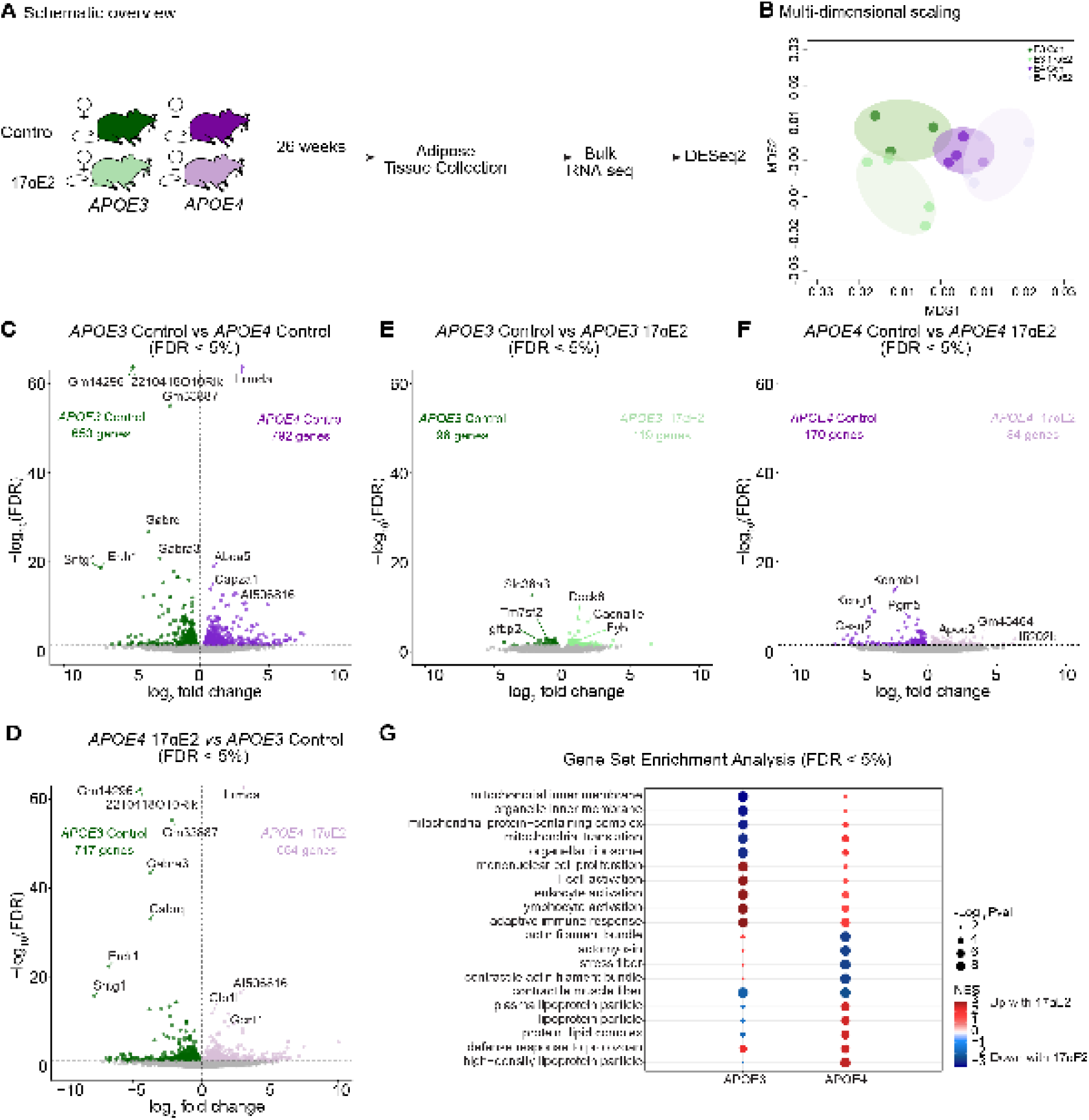
17αE2 differentially impacts lipoprotein processing in visceral adipose tissue of *APOE3* and *APOE4* mice. (A) Schematic overview. Visceral adipose tissue (VAT) depots were isolated from the mice after 26 weeks of treatment. (B) Multi-dimensional scaling (MDS) of transcriptomes for VAT from all four groups (n=3-4/group). (C) Differentially expressed genes between *APOE3* control and *APOE4* control VAT (FDR < 5%). (D) Differentially expressed genes between *APOE3* control and *APOE4* 17αE2 VAT (FDR < 5%). (E) Differentially expressed genes between *APOE3* control and *APOE3* 17αE2 (FDR < 5%). (F) Differentially expressed genes between *APOE4* control and *APOE4* 17αE2 (FDR < 5%). (G) Effect of 17αE2 in *APOE3* and *APOE4* VAT gene set enrichment analysis top 5 positive and negative NES found in both comparisons (DESeq2 FDR < 5%).

### 17αE2 improves metabolic outcomes in female APOE4 mice

Consistent with higher adiposity, *APOE4* control mice exhibited increased fasting glucose compared to *APOE3* control mice prior to treatment (**Figure 3B**). After 25 weeks of 17αE2 treatment, there was a significant effect of treatment with both *APOE3* and *APOE4* mice displaying reduced fasting glucose (**Figure 3B**). In the glucose tolerance test, *APOE4* control mice exhibited impaired glucose tolerance relative to *APOE3* control mice (**Figure 3C,D**), and 17αE2 treatment was associated with significant improvement only in *APOE4* mice (**Figure 3D**). Our previous work in middle-aged males showed *APOE4* associated with higher levels of hepatic steatosis compared to *APOE3* [43]. In females, while there was no significant genotype difference in hepatic steatosis as indicated by oil red O staining, there was a numerical trend for reduction by 17αE2 treatment only in *APOE4* mice (**Figure 3E**). Using a validated epigenetic clock for mouse aging, we found that *APOE4* animals had a liver DNA methylation signature consistent with that of chronologically older mice, while *APOE3* control and treated mice had a comparatively younger DNA methylation pattern. 17αE2 had no impact on DNAge in *APOE3* mice and trended toward increased DNAge in *APOE4* mice (**Figure 3F**). Together, these data identify increased metabolic impairment and aging phenotypes with the *APOE4* genotype in middle-aged female mice, but mixed effects of 17αE2 treatment.

**Figure 3.**
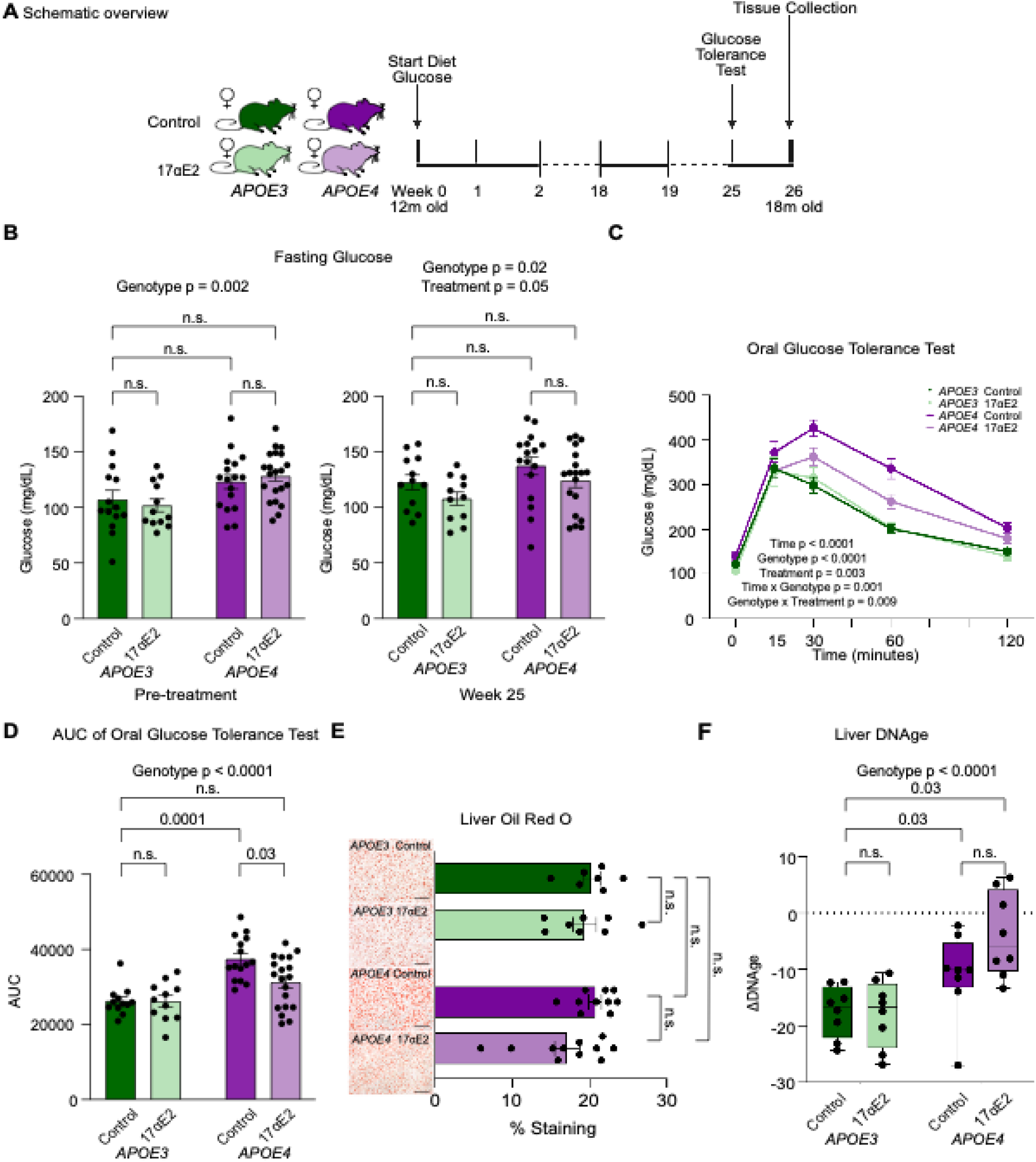
*APOE4* mice have greater metabolic dysfunction and greater protection with 17αE2 compared to *APOE3* mice. (A) Schematic overview. Glucose tolerance test was performed after 25 weeks of treatment and tissues were collected after 26 weeks. (B) Fasting glucose measured pre-treatment and after 25 weeks of 0 or 14.4 ppm 17αE2. (C) Oral glucose tolerance test (GTT) measured at baseline then 15-, 30-, 60-, and 120-minutes post oral gavage with glucose. (D) Area under the curve analysis for GTT seen in (C) (n=11-20/group). (E) ORO quantification (n=7-11/group). Images show oil red O labeling (ORO) of lipid accumulation in livers of 18-month-old female *APOE3* and *APOE4* mice treated with 0 or 14.4 ppm 17αE2. Scale bar size indicates 100µM. (F) Measure of biological aging using a DNA methylation clock derived from liver tissue (n=8/group). In (B), shaded panels indicate pre-treatment. In panel (C) dark green lines indicate *APOE3* control, light green lines indicate *APOE3* 17αE2, dark purple lines indicate *APOE4* control, and light purple lines indicate *APOE4* 17αE2. In (B) and (D-F) dark green indicates *APOE3* control, light green indicates *APOE3* 17αE2, dark purple indicates *APOE4* control, and light purple indicates *APOE4* 17αE2. Data show mean ± SEM.

To further investigate whether the metabolic improvements induced by 17αE2 treatment extend to aspects of energy expenditure, mice were placed into metabolic chambers at week 18 of treatment (**Figure 4A**). Upon entry into the metabolic cages, 17αE2 treated *APOE4* mice weighed significantly less than *APOE4* control mice (**Supplementary Figure 4A**). Heat production (energy expenditure) in mice is measured using indirect calorimetry, which is calculated using oxygen consumption (VO_2_) and carbon dioxide production (VCO_2_). There were no significant differences in energy expenditure (**Figure 4B**) or movement (**Supplementary Figure 4B**) by genotype, treatment, or body weight. In control mice, the respiratory exchange ratio (RER) trended to be lower in *APOE4* relative to *APOE3* mice, particularly in the dark time (**Figure 4C)**. Following 17αE2 treatment, RER was increased specifically in *APOE4* mice such that treated *APOE4* mice achieved levels comparable to the *APOE3* groups (**Figure 4C**). This is likely due to the decrease in body weight, with higher body weights correlating with lower RER in both *APOE3* and *APOE4* mice (**Figure 4D**). Thus, 17αE2 is associated with modest metabolic improvements in female mice with *APOE4,* with comparatively little to no effect in *APOE3* females.

**Figure 4.**
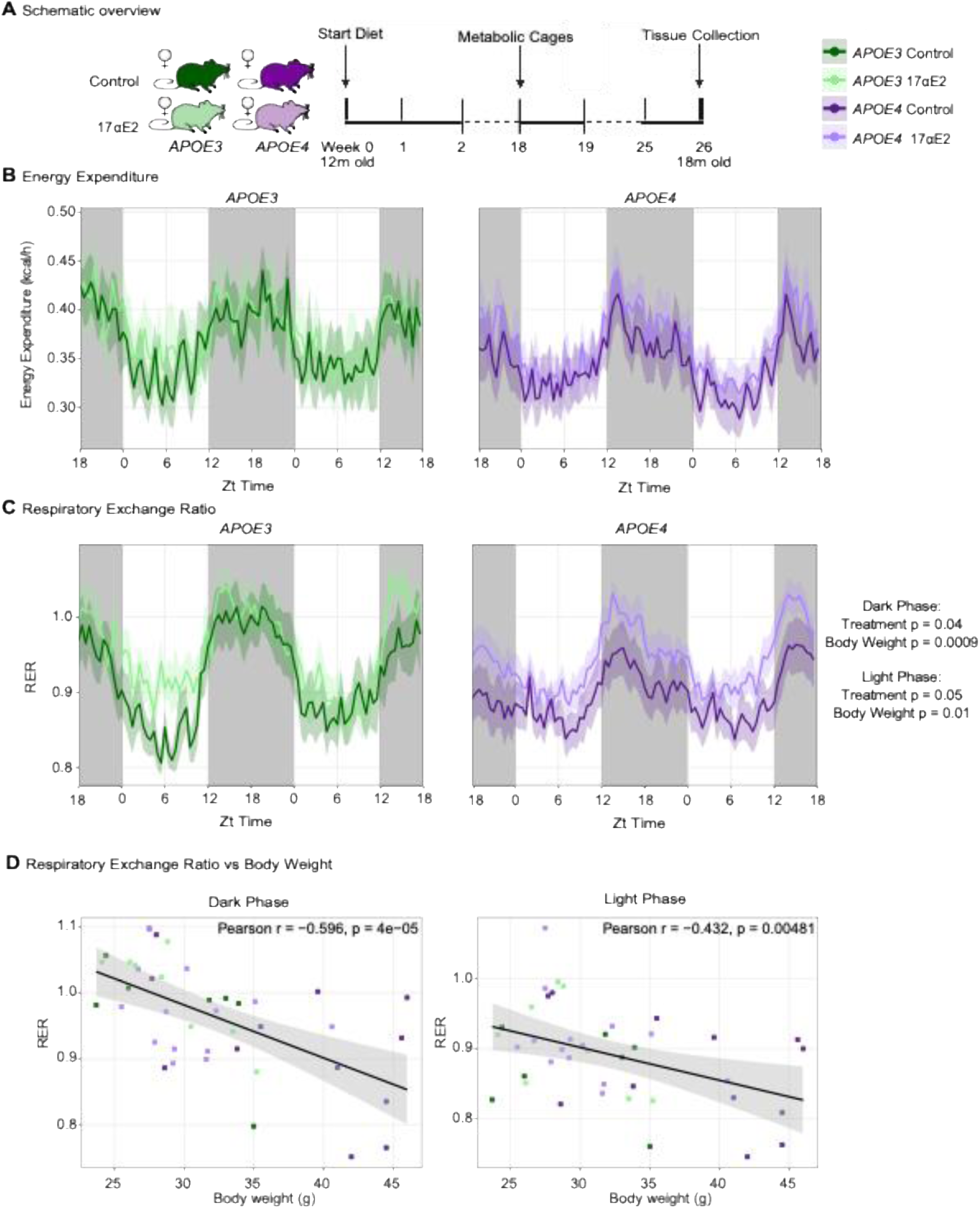
*APOE4* mice have increased respiratory exchange ratio with 17αE2 compared to control mice. (A) Schematic overview. Metabolic cage analysis was performed at 18 weeks of 17αE2 treatment. Data represents the last full 24 hours the mice were in the cages. (B) Heat expenditure measured through metabolic cages. (C) Respiratory exchange ratio calculated by VO2/VCO2. (D) Correlation between RER and body weight. For all measures shown n = 7-12/group. In panel (B-D) dark green lines indicate *APOE3* control, light green lines indicate *APOE3* 17αE2, dark purple lines indicate *APOE4* control, and light purple lines indicate *APOE4* 17αE2. Data show mean ± SEM.

### 17αE2 does not improve neural outcomes in APOE targeted replacement female mice

With 17αE2 treatment yielding metabolic benefits primarily in *APOE4* mice, we next asked if these improvements extend to the brain. At weeks 23 and 24 of treatment, mice were assessed for general motor and anxiety behaviors (open field), attention and working memory (spontaneous alternation behavior), recognition memory (novel object placement, recognition), and hippocampal-dependent spatial learning and memory (Barnes maze) (**Figure 5A**). In the open field test, there was a trend for increased time spent in the center with 17αE2 treatment, consistent with previous studies that found that the primary estrogen 17β-estradiol reduces anxiety in female rodents [44] (**Figure 5B**). Interestingly, *APOE4* mice traveled more distance irrespective of treatment, consistent with our previous study using *APOE* targeted replacement male mice [43] (**Figure 5C**). In the spontaneous alternation behavior test, there was a nonsignificant trend for *APOE4* control mice to perform worse than *APOE3* control mice, with no effect of 17αE2 treatment (**Supplementary Figure 5A**). In the Novel Object Placement (NOP) test, there is a weak trend for control *APOE4* mice to perform worse than *APOE3* mice of either treatment with a negative mean discrimination index indicating more time spent with the old object (**Figure 5D**). There was a non-significant trend (genotype x treatment p = 0.07) for NOP improvement in the 17αE2-treated *APOE4* group (**Figure 5D**). There were no significant genotype or treatment effects in Novel Object Recognition (**Supplementary Figure 5B**). In the Barnes maze, *APOE4* animals performed worse than *APOE3*, irrespective of treatment. This was consistent in multiple outcomes, including training errors (**Supplementary Figure 5C**), training path length to exit hole (**Figure 5E**), and training escape latency (**Figure 5F**). On the probe day, there was no improvement in time in the correct quadrant for either genotype (**Figure 5G**). Taken together, while 17αE2 had multiple peripheral improvements in the *APOE4* female mice, significant benefits were not seen behaviorally.

**Figure 5.**
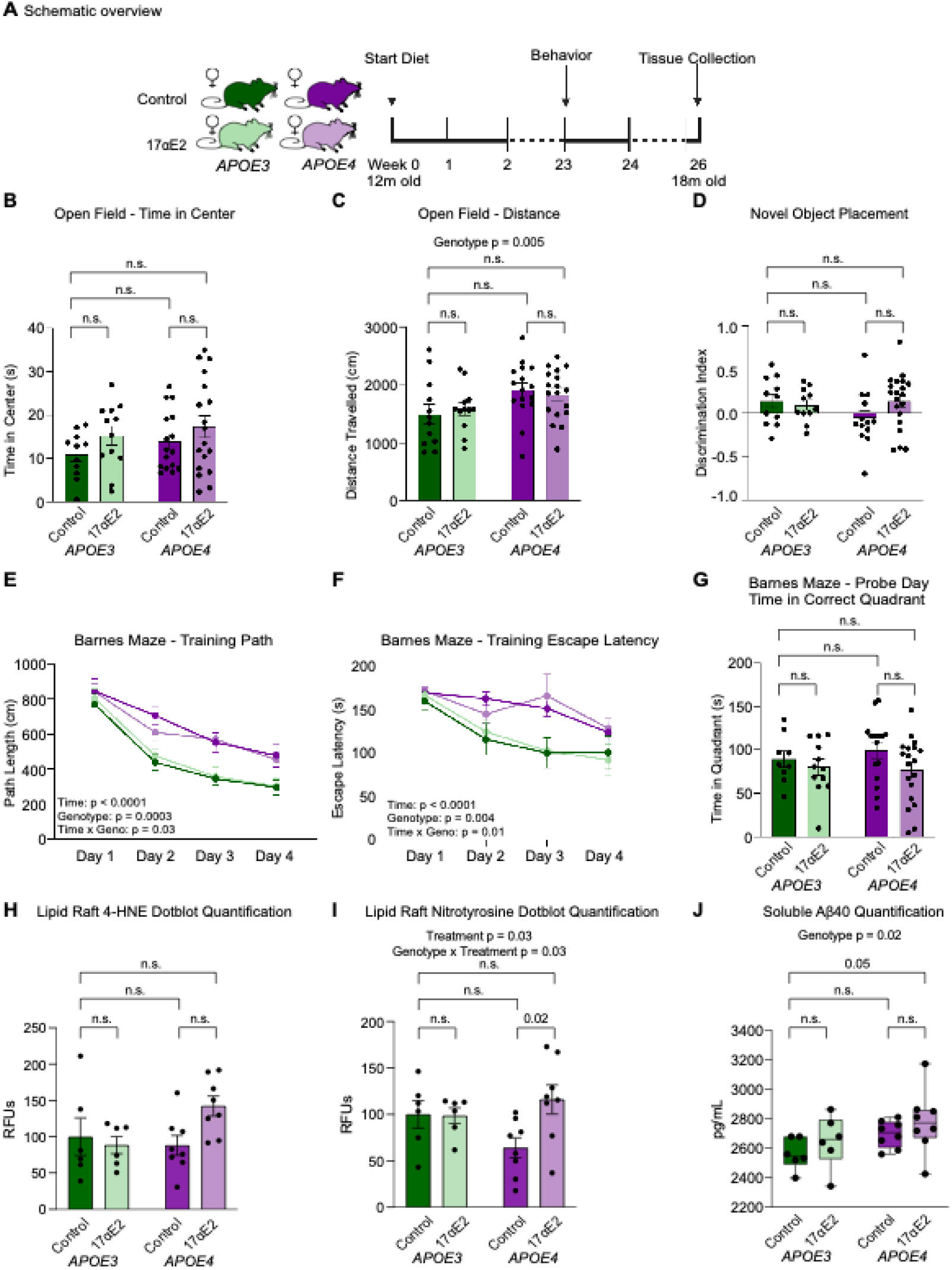
*APOE4* mice have greater neural deficits than *APOE3* mice. (A) Schematic overview. Animals were subjected to behavior tests after 23 weeks of treatment. (B) Time in center in the open field task (n=11-18/group). (C) Distance travelled in the open field task (n=11-18/group). (D) Novel object placement discrimination index. Positive values indicate more time spent with novel object; negative values indicate more time spent with old object (n=11-18/group). (E) Total path length to exit hole during the Barnes maze training days (n=11-18/group). (F) Escape latency during the Barnes maze training days (n=11-18/group). (G) Total time in correct quadrant during the Barnes maze probe trial (n=11-18/group). (H) Quantification of the lipid peroxidation marker 4-hydroxynonenal (4-HNE) from cortical lipid rafts (n=6-8 mice/group). (I) Quantification of the oxidative damage marker nitrotyrosine from cortical lipid rafts and representative dot blot images (n=6-8 mice/ group). (J) Quantification of soluble b-amyloid peptide (Ab-40) from Meso Scale Discovery plate (n=6-8 mice/ group). In (B) through (D) and (H) through (J), dark green indicates *APOE3* control, light green indicates *APOE3* 17αE2, dark purple indicates *APOE4* control, and light purple indicates *APOE4* 17αE2. In (E) and (F), dark green lines indicate *APOE3* control, light green lines indicate *APOE3* 17αE2, dark purple lines indicate *APOE4* control, and light purple lines indicate *APOE4* 17αE2. Data show mean ± SEM.

Next, we examined biochemical indices associated with brain aging, *APOE* genotype, and AD. First, we assessed brain oxidative damage by measuring 4-hydroxynonenal (4HNE), a lipid peroxidation byproduct and established biomarker of oxidative stress[45]. Lipid peroxidation increases during normal brain aging and is further elevated in AD[45–48], particularly within lipid rafts. In male mice, we previously reported higher 4HNE in cerebrocortical lipid rafts from *APOE4* control mice relative to the *APOE3* controls, with a reduction by 17αE2 treatment only in *APOE4* mice [25]. In female *APOE* mice, there was no significant difference in 4HNE by *APOE* genotype in control mice; however, 17αE2 treatment increased 4HNE only in the *APOE4* group (**Figure 5H**). We also measured nitrotyrosine, another marker of oxidative damage, and again found that the only significant effect was increased levels in the *APOE4* 17αE2-treated group (**Figure 5I**). Next, we measured soluble levels of amyloid-beta (Ab), a protein implicated in AD pathogenesis, in cerebral cortex homogenates. In males, we reported that 17αE2 treatment reduced cerebrocortical Abx-40 in *APOE4* but not *APOE3* mice [25]. In females, while there were significantly higher Abx-40 levels in *APOE4* control than *APOE3* mice, there was no significant effect of treatment (**Figure 5J**). There were no significant differences by genotype or treatment in soluble levels of Aβx-38 or Aβx-42 peptides (**Supplementary Figure 5D,E**). Taken together, *APOE4* females treated with 17αE2 displayed increased markers of oxidative damage without changes in soluble Ab levels.

## Discussion

Late-onset Alzheimer’s disease risk is strongly influenced by the *APOE4* genotype, female sex, and age. While many longevity interventions exhibit sex-specific effects, there is a limited understanding of how these interventions impact healthspan in a sex-dependent manner. In this study, we explored the effects of the male-longevity-promoting compound 17αE2 on female healthspan within the *APOE* genotype context. Specifically, we assessed the ability of 17αE2 to protect against *APOE4*- and age-associated metabolic and behavioral impairments in female mice carrying human *APOE3* or *APOE4* targeted gene replacement. Prior work from our group demonstrated that 17αE2 had genotype-specific effects, primarily benefiting *APOE4* males [25], while other studies noted limited effects in females expressing murine Apoe [19, 49].

In the present study, 17αE2 was administered to female mice at 12 months of age, after the establishment of genotype-specific metabolic and behavioral differences and continued for 6 months. Our findings revealed that 17αE2 mitigated metabolic impairments in *APOE4* females but showed limited benefit on neural outcomes. Metabolic improvements, including reduced body weight, increased lean mass, improved glucose tolerance, and higher energy expenditure, were observed only in *APOE4* females. This contrasts with our prior work in males, where 17αE2 conferred improvements in both *APOE3* and *APOE4* mice, with larger effect sizes in *APOE4* males. Transcriptomic analysis of visceral adipose tissue revealed enrichment of high-density lipoprotein particle-related gene sets in *APOE4* mice treated with 17αE2, while these pathways were suppressed in treated *APOE3* mice. This divergence may result from the structural and functional differences between the *APOE* isoforms. *APOE4’s* single amino acid substitution (Arg112 instead of Cys112) destabilizes its structure and reduces lipid-binding capacity[5, 6]. Indeed, higher HDL trafficking enhances the transport and recycling of cholesterol and other lipids to and from peripheral tissues, promoting reverse cholesterol transport back to the liver for excretion [50]. This process supports cardiovascular health by reducing cholesterol accumulation in arterial walls, lowering the risk of atherosclerosis, and maintaining cellular lipid homeostasis [50]. Estrogen influences lipid processing by enhancing the expression and activity of enzymes regulating lipid metabolism and apolipoprotein function, potentially alleviating some detrimental effects of *APOE4*[51, 52]. The pronounced differences in gene sets related to lipoprotein processing may pose a potential mechanism by which 17αE2 improves *APOE4* systemic outcomes.

In neural assessments, both genotypes showed reduced anxiety with 17αE2, consistent with prior studies on estrogens [44], but no rescue in spatial learning deficits was noted for *APOE4* mice. *APOE4* females exhibited equal levels of oxidative damage markers compared to *APOE3*, and 17αE2 increased these markers with no effect on amyloid. In *APOE4* males, 17αE2 reduced lipid raft–associated oxidative damage and lowered cerebrocortical Aβx-40, consistent with a neuroprotective profile. In the DNAge assessment, 17αE2 treatment increased biological age in the female liver, opposite to the effect seen in males [25]. These mixed effects could be due to the presence of a ‘critical window’ in the timing of estrogenic treatment. The concept of a critical window for hormone replacement therapy has been extensively explored in the context of female aging and menopause [27, 53]. Although mice do not experience menopause, they undergo estropause by 12 months of age [54]. Female mice have significantly reduced primordial follicles with age, with at least a 50% reduction seen at 10 months and a 74% reduction by 12 months in C57BL/6 mice [55, 56]. Thus, the timing of 17αE2 administration in our study coincides with a period of declining endogenous estrogen signaling. *APOE4* females displayed uterotrophic responses to 17αE2, similar to that seen in ovariectomized middle-aged UM-HET3 mice[22], while *APOE3* females did not, highlighting genotype-specific differences in estrogen responsiveness. One possible explanation is that the 17αE2 dose used in this study is sufficient to elicit estrogenic effects in *APOE4* females but falls below the threshold required to induce similar responses in *APOE3* females. Alternatively, *APOE3* females may maintain endogenous estrogen signaling for a longer period of time, reducing their physiological responsiveness to exogenous estrogenic compounds at this age. Together, these observations point to genotype-dependent differences in estrogen sensitivity that may influence how female animals respond to hormone-related interventions later in life.

Estrogen depletion during menopause is associated with increased adiposity, metabolic dysregulation, and cognitive decline in humans [57–60]. Clinical studies indicate that 17β-estradiol (17βE2) replacement post-menopause reduces adiposity and overall metabolic risk, though the timing of therapy significantly influences its efficacy[57, 61–64]. The literature surrounding the role of estrogen replacement in reducing AD risk factors is inconsistent, with some reports showing neutral or adverse effects if treatment begins after menopause [65]. Animal studies show 17βE2 can confer significant metabolic and cognitive benefits when administered later in life [66, 67], but results are conflicting [68, 69]. *APOE* genotype adds further complexity, as responses to estrogen therapy in the *APOE4* context are conflicting, with both improved [43, 70] and diminished [71, 72] outcomes reported.

These results add to the growing literature defining the impact of aging and *APOE* genotype on the efficacy of estrogen-based therapies in protecting against AD and other dementias. This study highlights the genotype-specific effects of 17αE2 on metabolic and behavioral healthspan in female mice. While *APOE4* females exhibited significant metabolic benefits from 17αE2 treatment, including improved glucose tolerance and energy expenditure, behavioral improvements were more limited, and *APOE3* females demonstrated minimal responsiveness to the intervention. Notably, the lack of significant lifespan extension in previous studies of 17αE2 does not preclude its ability to promote healthspan, as evidenced by the metabolic benefits observed in *APOE4* mice. This emphasizes the need to separate the evaluation of lifespan and healthspan outcomes in aging research. These findings also highlight the importance of incorporating females into studies of aging interventions. As a follow up to our previous study in males [25], this study addresses this gap and provides valuable insights into how sex and genotype influence 17αE2’s efficacy. Future investigations should prioritize earlier intervention time points before the onset of estrogen depletion to better understand the preventive potential of 17αE2 and other therapies in addressing both metabolic and cognitive aspects of aging. Additionally, exploring the molecular pathways underlying the differential responses between *APOE* genotypes may reveal novel targets for mitigating the adverse effects of aging and *APOE4*-driven risks in women.

## Limitations to the study

There are a few limitations to this study. First, genetic background can influence the efficacy of longevity-promoting interventions [73, 74], and while foundational studies of 17αE2 longevity effects were conducted in UM-HET3 mice [19, 22], the present work used mice on a predominantly C57BL/6 background. Thus, strain-specific factors may have shaped the responses observed in *APOE3* and *APOE4* humanized mice. Second, *APOE* allele zygosity may modulate responsiveness to 17αE2. Our recent work demonstrates that *APOE3/4* heterozygous mice exhibit intermediate or distinct metabolic, neuroimmune, and behavioral phenotypes compared to *APOE3/3* and *APOE4/4* mice, indicating that zygosity-dependent estrogen responsiveness warrants further investigation [75]. Third, although 18-month-old mice exhibit cognitive deficits and neuroinflammation in this model, studies in older animals may better capture advanced age-related neural impairments. Finally, while our findings suggest a potential protective role for 17αE2 against APOE4-associated cognitive decline, this study does not directly interrogate AD pathology. Future work examining 17αE2 in models incorporating amyloidosis and tauopathy will be essential to fully assess its relevance to AD-related mechanisms.

## Data availability

RNA-seq analyses data is available in **Supplementary Table 1**. All sequencing data were deposited to SRA under accession PRJNA1425111. DNAge data were deposited to FigShare (doi:10.6084/m9.figshare.31558918). All other data are available from the corresponding author upon reasonable request.

## Author Contributions

C.J.P., B.A.B., and C.E.F. designed the study. C.J.M, A.C., S.N., H.A., M.V., and M.A.T. performed experiments. C.J.M. performed data analyses and bioinformatics analyses. C.J.M., B.A.B. and C.J.P. wrote the manuscript. All authors approved the final version of the manuscript.

## Acknowledgements

This study was supported by a grant from the Cure Alzheimer’s Fund (C.J.P., C.E.F., B.A.B.). C.J.M. was supported by NIH/NIA grants T32 AG052374 (S.P. Curran), F31 AG084279 (C.J.M.), and T32 AG000213 (R. Anderson).

