## Supplementary Figure for "17α-Estradiol Confers Limited Protection Against *APOE4* Phenotypes in Middle-Aged Female Mice"

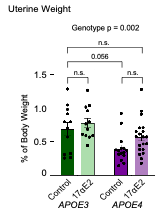

**Supplementary Figure 1. *APOE4* uterine weight increases with 17αE2 treatment.** (A) Percent of dry uterine weight compared to overall body weight after 26 weeks of 0 or 14.4ppm 17αE2 (n=11-20/group). In (A), dark green indicates *APOE3* control, light green indicates *APOE3* 17αE2, dark purple indicates *APOE4* control, and light purple indicates *APOE4* 17αE2. Data show mean ± SEM. Asterisks denote statistical significance: * p < 0.05 in 2-way ANOVA Tukey post-hoc test.

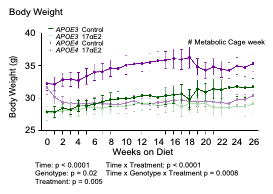

**Supplementary Figure 2. Adiposity is decreased with 17αE2 treatment.** (A) Body weights of *APOE3* and *APOE4* female mice throughout 26 weeks of treatment (n=11-20/group). In (A), dark green indicates *APOE3* control, light green indicates *APOE3* 17αE2, dark purple indicates *APOE4* control, and light purple indicates *APOE4* 17αE2. Data show mean ± SEM.

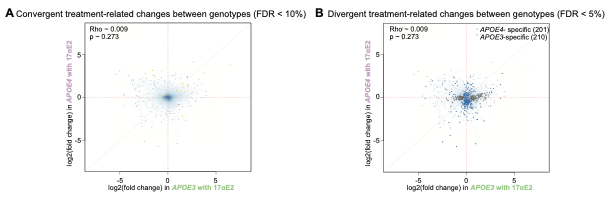

**Supplementary Figure 3. Convergent and divergent treatment-related changes between genotypes.** (A) Convergent treatment-related changes in the effect of 17αE2 between genotypes (FDR < 10%). (B) Divergent treatment-related changes in the effect of 17αE2 between genotypes (FDR < 5%).

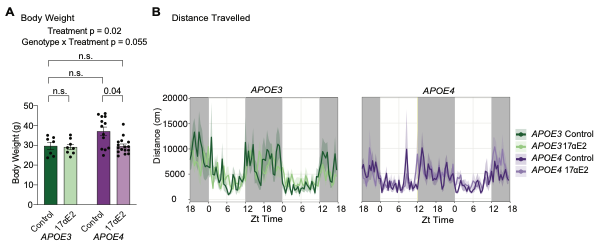

**Supplementary Figure 4. Metabolic cage body weight and distance travelled.** (A) Body weight of metabolic cage mice at week 18 of the study. (B) Distance traveled (cm) over the course of 48 hours in the metabolic cages. (In (A) and (B), dark green indicates *APOE3* control, light green indicates *APOE3* 17αE2, dark purple indicates *APOE4* control, and light purple indicates *APOE4* 17αE2. For all measures shown n = 7-12/group. Data show mean ± SEM.

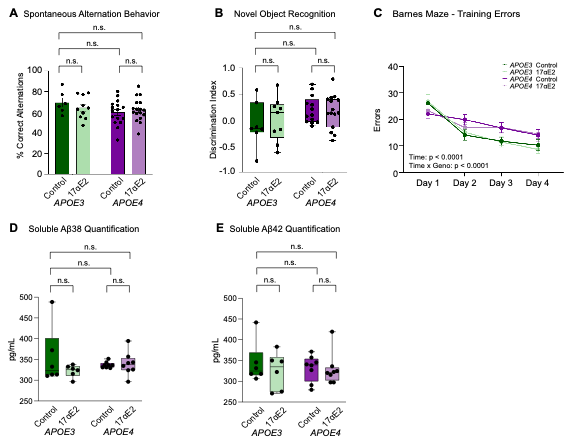

**Supplementary Figure 5. Working memory and short-term hippocampal memory are unchanged with 17αE2 treatment.** (A) Spontaneous alternation behavior presented as % of correct alternations (n=6-16/group). (B) Novel object recognition discrimination index. Positive values indicate more time spent with novel object; negative values indicate more time spent with old object (n=7-15/group). (C) Total errors during the Barnes maze training days (n=11-18/group). (D) Quantification of soluble b-amyloid peptide (Ab-38) from Meso Scale Discovery plate (n=6-8 mice/ group). (E) Quantification of soluble b-amyloid peptide (Ab-42) from Meso Scale Discovery plate (n=6-8 mice/ group). In (A) through (E), dark green indicates *APOE3* control, light green indicates *APOE3* 17αE2, dark purple indicates *APOE4* control, and light purple indicates *APOE4* 17αE2. Data show mean ± SEM.
